# Alisertib exerts KRAS allele specific anticancer effects in colorectal cancer

**DOI:** 10.1101/2020.10.14.340125

**Authors:** Baojun Ren, Zhuowei Gao, Kehong Zheng, Yong Yang, Pengfei Su, Qimei Luo, Jing Feng, Zhentao Luo, Yan Geng, Yongle Ju, Zonghai Huang

## Abstract

Colorectal cancer (CRC) is a common cancer causing substantial mortality and morbidity worldwide. Oncogene *RAS* mutations occur notably in ∼45% of CRCs, associated with a poor prognosis. KRAS is subject to multiple tiers of regulation, including kinase. Aurora kinases has been implicated in many types of tumor onsets and progression, making them as a promising therapeutic targets. Alisertib (ALS), selectively inhibits Aurora kinase A (AURKA) and exerts potent anticancer activities *in vitro* and *in vivo* studies, but the latent anticancer effect of ALS on CRC remains unclear in the context of different KRAS mutations. This study aimed to assess the effects of ALS on RAS signaling pathway in a panel of CRC lines expressing different KRAS alleles, including Caco-2 (KRAS WT), Colo-678 (KRAS G12D), SK-CO-1 (KRAS G12V), HCT116 (KRAS G13D), CCCL-18 (KRAS A146T), and HT29 (BRAF V600E). The results showed that ALS modulated the active form of KRAS in a RAS allele specific manner across the panel of CRC lines; ALS differentially regulated RAS signal via PI3K/Akt and MAPK pathways; and ALS induced apoptosis and autophagy in a RAS allele specific manner. Of note, in combination of ALS and MEK inhibitor, selumetinib, enhanced ALS regulatory effects in CRC lines in a RAS allele specific manner on apoptosis, autophagy, and cell growth. Taken together, this study suggests that ALS differentially regulates RAS signaling pathway and manipulates cell apoptosis and autophagy in RAS allele specific manner. The combinatorial approach of ALS and MEK inhibitor may represent a new therapeutic strategy for precision therapy of CRC in a RAS allele manner.

## Introduction

Colorectal cancer (CRC) is one of the top 3 common cancers worldwide and causes high mortality and morbidity [1]. An estimation of 147,950 new cases and 53,200 deaths occurred in United States, rendering CRC as one of the top 3 common cause of cancer related death in 2020 [2]. It was one of the top 5 common cancer and one of the common cause of cancer death in 2011 in China [3]. There were 253,000 new CRC cases, 139,000 cancer deaths, and 583,000 people living with cancer (within 5 years of diagnosis) in 2012 in China [1]. Notably, these numbers are increasing. The overall 5-year survival rate is lower but the metastatic CRC patient number is higher relative to that in United States [4, 5]. Unfortunately, an estimation of 20%-30% of patients were newly diagnosed with unresectable metastatic CRC and 50%-60% patients will develop metastatic CRC [6], which substantially jeopardizes the therapeutic outcome in clinic. Thusly, it urgently requires the exploration of new therapeutic strategy for CRC treatment.

RAS mutations are found in a large proportion of deadly human tumors, occurring in 20-30% of malignant tumors overall, and in about 45% of CRCs. RAS mutations are essential for tumor initiation and maintenance [7, 8]. Moreover, RAS mutations are often associated with a poor prognosis [9-11]. RAS proteins are guanosine triphosphatases (GTPases) which function as binary switches that cycle between in inactive (GDP bound) and active (GTP bound) states [12, 13]. Activated RAS proteins bind to numerous effector proteins including RAF kinases, PI3K, RalGDS among others which regulate critical cellular processes including metabolism, proliferation, and survival [14]. Conversion from GDP bound RAS to GTP is stimulated by guanine nucleotide exchange factors (GEFs) while conversion back to the inactive GDP bound form is mediated by GTPase activating proteins (GAPs) [12, 15]. Critical to the activation of RAS proteins is membrane localization mediated prenylation of the C-terminal end [16]. Somatic mutations in RAS proteins alter the normally tightly regulated process of RAS signaling by causing an accumulation of GTP-bound RAS.

Aurora kinase A, B, and C (AURKA/B/C) are the three Aurora kinase members that play a crucial role in regulating cell mitosis, including centrosome duplication, spindle assembly, chromosome alignment, chromosome segregation, and the fidelity-monitoring spindle checkpoint [17, 18]. A large body of evidence shows a correlation between the aberration of AURKA expression at transcriptional and translational levels and cancer development, including breast, pancreatic, ovarian and gastric cancers [19], which can be ascribed at least to the development of aneuploidy, supernumerary centrosomes, defective mitotic spindles, and resistance to apoptosis. Of note, an association between the aberrant expression of AURKA and poor prognosis and response to chemotherapy in CRC was observed [20, 21]. For example, in relative to the primary tumor, a higher expression of AURKA was detected in CRC liver metastasis which can be a molecular biomarker, independent of established clinicopathological variables [22]. Also, the essentiality of AURKA for CRC stem cells regeneration and resistance to cytotoxic stimuli was observed [23].

Previously, we assessed the regulatory effects of alisertib (ALS), a selective inhibitor of AURKA, on cell growth, migration, apoptosis, and autophagy in wild type KRAS and BRAF V600E mutant CRC cell lines. However, the comparison on the effects of AURKA inhibition was not determined in the context of different common RAS mutational background, including KRAS G12D, G12V, G13D, and A146T. In this study, we aimed to assess the differential regulatory effects of ALS on PI3K/Akt and MAPK signaling pathways and cell apoptosis, autophagy, and cell growth in a panel of human CRC cell lines with different RAS mutations and engineered Flp-In T-REx cells.

## Materials and methods

### Chemicals and reagents

ALS and Selumetinib (Sel) were bought from Selleckchem Inc. (Houston, TX, USA) and stored at 100 mM in Dimethyl sulfoxide (DMSO) at −20°C. DMSO, fetal bovine serum (FBS), ammonium persulfate, protease and phosphatase inhibitor cocktails, doxycycline (DOX), and Dulbecco’s phosphate buffered saline (PBS) were purchased from Sigma-Aldrich (St Louis, MO, USA). All required medium including EMEM, McCoy’s 5A, RPMI1640, and DMEM were obtained from Corning Cellgro Inc. (Herndon, VA, USA). Pierce™ bicinchoninic acid (BCA) protein assay kit and radioimmunoprecipitation assay (RIPA) buffer were sourced from Thermo Fisher Scientific Inc. (Waltham, MA, USA). Western blotting substrate was bought from Cell Signaling Technology (Beverly, MA, USA). Skim milk and nitrocellulose membrane were bought from BioRad (Hercules, CA, USA). Cleaved PARP, p-Akt (Ser473), Akt, p-Erk1/2 (Thr202/Tyr204), Erk1/2, p-GSK3α/β (Ser21/9), RAS, and LC3-I/II antibodies were all purchased from Cell Signaling Technology Inc. (Beverly, MA, USA), and β-actin was bought from Santa Cruz Biotechnology Inc. (Dallas, TX, USA).

### Cell lines

Caco-2 (KRAS WT), Colo-678 (KRAS G12D), SK-CO-1 (KRAS G12V), HCT116 (KRAS G13D), CCCL-18 (KRAS A146T), and HT29 (BRAF V600E) are well-recognized colorectal cancer cell lines and were bought from ATCC (Manassas, VA, USA). Cells were cultured in ATCC recommended complete medium and maintained in a 5% CO_2_/95% air humidified incubator at 37°C. Flp-In T-REx 293 cells expressing KRAS WT and G12D were generated by following manufacturer’s instructions (Thermal Fisher). Mycoplasma Plus PCR Primer Set (Agilent) was used to determine mycoplasma for all cell lines. Cells were treated with ASL or Sel with 0.05% DMSO.

### Cell growth assessment

To assess the effects of inhibitors of AURKA and MEK on cell growth, engineered KRAS WT and G12D expressing Flp-In T-REx 293 cells (1 × 10^3^) were utilized in the presence of DOX (2 ng) upon the exposure of ALS and Sel. Cell growth was monitored in IncuCyte and images were taken every 4 hrs in three fields per well in black 96-well plates with clear bottom. Data were analyzed and plotted by PRISM.

### Ras-GTP Pull-down

Colorectal cancer cell lines (Caco-2, Colo-678, SK-CO-1, HCT116, CCCL-18, and HT29) were grown in 10% FBS in 60 mm dishes and treated with ALS for 48 hrs. Proteins samples were processed in SDS free lysis buffer for the assessment of Ras-GTP level using Active Ras Detection Kit (Cell Signaling) by following manufacturer’s instructions.

### Western blotting assay

The effects of ALS and Sel on the protein expression level and RAS signaling pathway were examined using Western blotting assays. Colorectal cancer cell lines (Caco-2, Colo-678, SK-CO-1, HCT116, CCCL-18, and HT29) were collected and protein samples were processed in RIPA buffer containing phosphatase and protease inhibitor cocktails after 48-hr treatment with inhibitors, then the samples were subject to Western blotting assay.

### Statistical analysis

Data are presented as the mean ± standard deviation (SD). Multiple comparisons were assessed by one-way analysis of variance (ANOVA) followed by Tukey’s multiple comparison procedure. Values of *P*<0.05 were considered statistically significant. All the assays were performed in triplicate.

## Results

### ALS modulates active form of RAS in a RAS allele specific manner

In order to test the effect of inhibition of AURKA on RAS signaling output, we first examined the level of RAS-GTP in a panel of CRC cell lines when treated with ALS. We used the RAF1-RBD to pull down the active form of RAS in Caco-2 (KRAS WT), Colo-678 (KRAS G12D), SK-CO-1 (KRAS G12V), HCT116 (KRAS G13D), CCCL-18 (KRAS A146T), and HT29 (BRAF V600E) cells. As shown in Figure 1, in wild-type KRAS expressing Caco-2 cells, ALS concentration-dependently increased the level of RAS-GTP, but there was no alteration in Colo-678 cells expressing KRAS G12D. ALS increased RAS-GTP level in SK-CO-1 cells expressing KRAS G12V and CCCL-18 cells expressing KRAS A146T, whereas there was no change in HCT116 cells expressing KRAS G13D. Of note, in HT29 cells expressing BRAF V600E, which is a common in CRC and constitutively active BRAF mutant and does not rely on RAS activation to activate RAS-RAF-MEK-ERK signal, ALS decreased the active form of RAS. Together, these results suggests that inhibition of AURKA has regulatory effect on RAS signaling pathway in a RAS allele specific manner.

**Figure 1.**
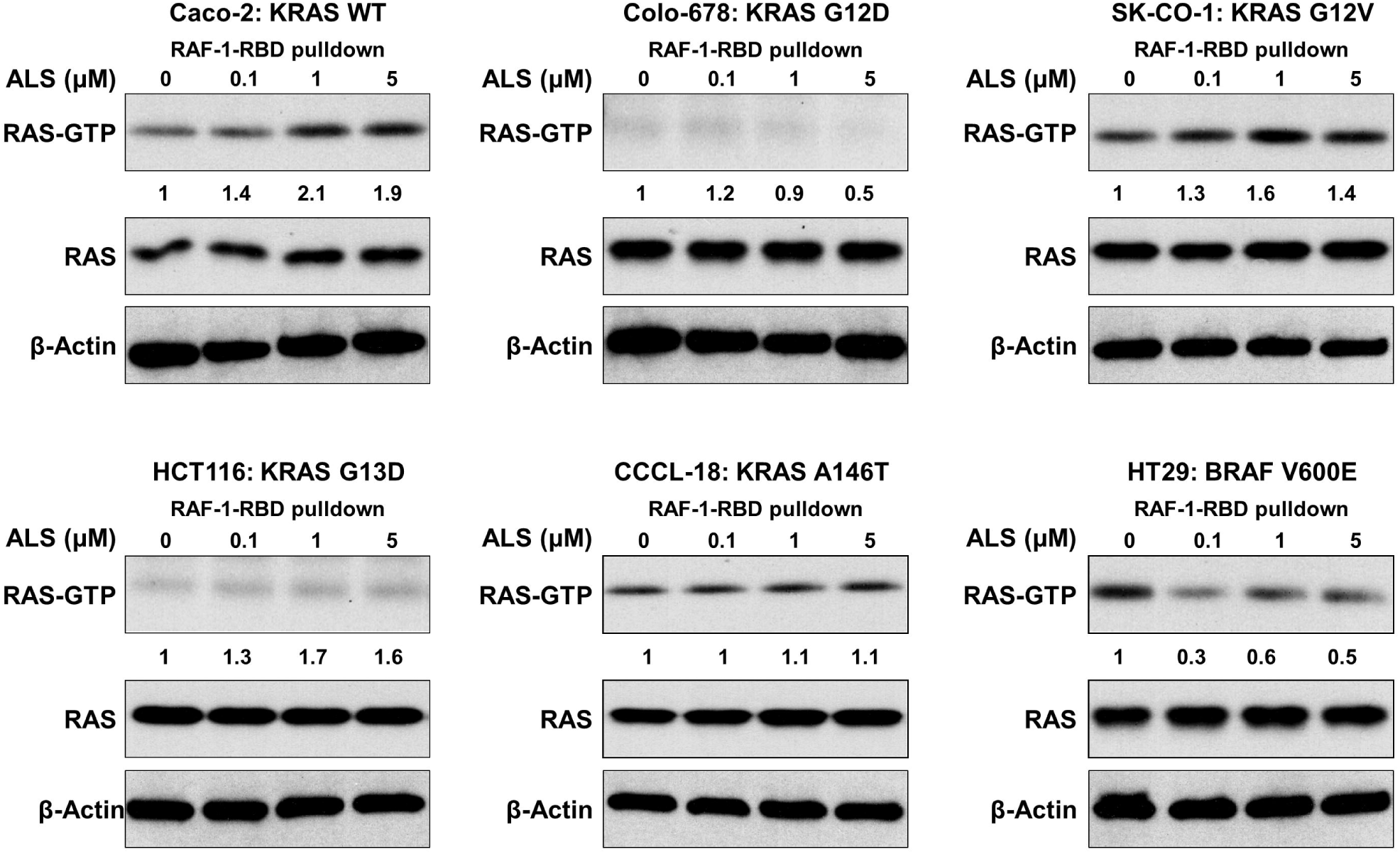
ALS modulates RAS-GTP level in a RAS allele specific manner in CRC cell lines. A panel of CRC cell lines including Caco-2 (KRAS WT), Colo-678 (KRAS G12D), SK-CO-1 (KRAS G12V), HCT116 (KRAS G13D), CCCL-18 (KRAS A146T), and HT29 (BRAF V600E) were treated with ALS and proteins were subject to pulldown using RAF-RBD and Western blotting assay for RAS and β-actin.

### ALS affects RAS signaling in a RAS allele specific manner

Following the examination of RAS-GTP level, we tested the phosphorylation level of Akt and Erk in the CRC cell lines. PI3K/Akt and MAPK signaling pathways are the two dominant downstream pathways of RAS. Upon the activation of RAS, the phosphorylation level of Akt and Erk are major readout. We treated the cells with ALS at 0.1, 1, and 5 µM and examined p-Akt and p-Erk. As shown in Figure 2, ALS inhibited the phosphorylation of Akt but promoted the phosphorylation of Erk in wild-type KRAS expressing Caco-2 cells. There was no change in the level of p-Akt and p-Erk in KRAS G12D expressing Colo-678 cells and KRAS G12V expressing SK-CO-1 cells. In KRAS G13D expressing HCT116 cells, ALS suppressed the phosphorylation of Akt and Erk. Similarly, ALS repressed the activation of Akt and Erk in BRAF V600E expressing HT29 cells. However, ALS enhanced the phosphorylation of Akt and Erk in CCCL-18 cells expressing KRAS A146T. Taken together, these results indicates that inhibition of AURKA via ALS leads to RAS allele specific modulation of PI3K/Akt and MAPK signaling pathway.

**Figure 2.**
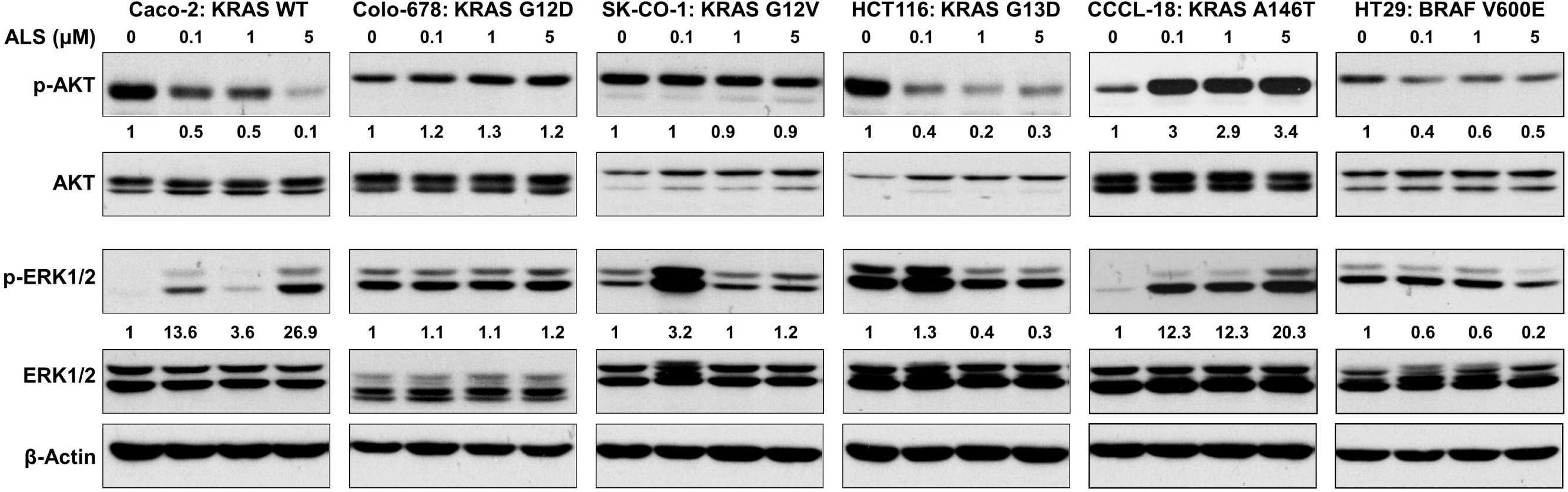
Inhibition of AURKA affects PI3K/AKT and MAPK signaling in a RAS allele specific manner in CRC cell lines. A panel of CRC cell lines including Caco-2 (KRAS WT), Colo-678 (KRAS G12D), SK-CO-1 (KRAS G12V), HCT116 (KRAS G13D), CCCL-18 (KRAS A146T), and HT29 (BRAF V600E) were treated with ALS and proteins were subject to Western blotting assay for assessment of phosphorylation of ERK and AKT.

### ALS manipulates cell apoptosis and autophagy in a RAS allele specific manner

To test the influence of RAS allele specific regulatory effect of ALS on cell death, we examined the level of cleaved-PARP and LC3I/II as a surrogate marker of cell apoptosis and autophagy. The panel of CRC cell lines were treated with ALS and the level of cleaved PARP and LC3I/II was measured. As shown in Figure 3, Wild-type KRAS expressing Caco-2 cells underwent apoptosis and autophagy upon the exposure of ALS, evident from the increase in the level of cleaved-PARP and ratio of LC3II/I. There was no marked change of apoptosis and autophagy in KRAS G12D expressing Colo-678 cells when treated with ALS. However, there was a striking increase in the level of cleaved-PARP and ratio of LC3II/I in SK-CO-1 cells expressing KRAS G12V. In addition, in HCT116, CCCL-18 and HT29 cells, ALS markedly induced the cleavage of PARP, to a lesser extent, ALS also increase the ratio of LC3II/I in CCCL-18 and HT29 cells. Collectively, pharmacological inhibition of AURAK via ALS results in different regulatory effect on cell apoptosis and autophagy in a RAS allele specific manner.

**Figure 3.**
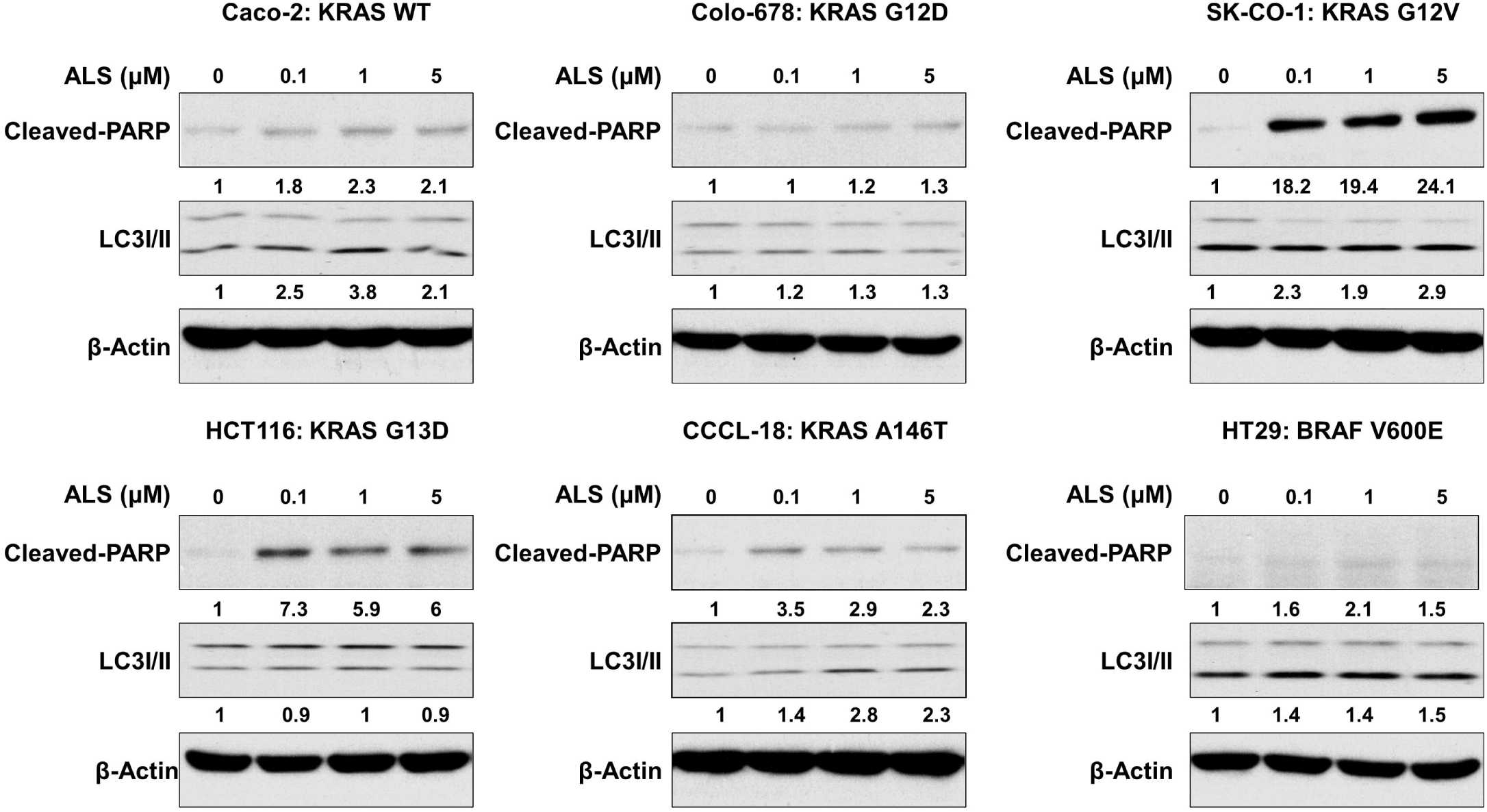
ALS manipulates cell apoptosis and autophagy in a RAS allele specific manner in CRC cell lines. A panel of CRC cell lines including Caco-2 (KRAS WT), Colo-678 (KRAS G12D), SK-CO-1 (KRAS G12V), HCT116 (KRAS G13D), CCCL-18 (KRAS A146T), and HT29 (BRAF V600E) were treated with ALS and cell apoptosis and autophagy were examined as indicated by cleaved-PRAP and LC3 by Western blotting assay.

### ALS combines MEK inhibitor resulting in an enhanced regulatory effects on RAS signals in a RAS allele specific manner

Since we found the RAS allele specific regulatory effects on RAS signaling pathways of ALS in CRC cell lines, we further asked if a combinatorial approach of ALS and MEK inhibitor enhanced ALS’s effects. To test the anticancer effects of pharmacological inhibition of AURKA and MEK on different RAS expressing background, we first tested the effects of MEK inhibitor, selumetinib (Sel) on PI3K/Akt and MAPK signaling pathways. As shown in Figure 4, Sel inhibited MAPK signaling pathway in all tested cell lines, evident from the suppression of phosphorylation of Erk, whereas, PI3K/Akt signaling pathway showed a differentiate responses to Sel treatment. p-Akt level was decreased in Caco-2, Colo-678 and SK-CO-1 cells, but increased in the rest cell lines. Furthermore, we tested the effect of Sel on cell apoptosis and autophagy across the panel of CRC cell lines. As shown in Figure 5, Sel did not affect cell apoptosis in Caco-2, Colo-678, and CCCL-18 cells, evident from the unaffected the cleavage of PARP. On the other hand, Sel induced cell apoptosis in SK-CO-1, HCT116, and HT29 cells, indicated by the increase in the level of cleaved-PARP. Also, Sel promoted cell autophagy via increasing the conversion of LC3I to LC3II in Caco-2, HCT116, CCCL-18, and HT29 cells. In aggregate, the results suggest that MEK inhibition via Sel results in differentiate responses in RAS signaling pathways, resulted in differential responses in cell apoptosis and autophagy.

**Figure 4.**
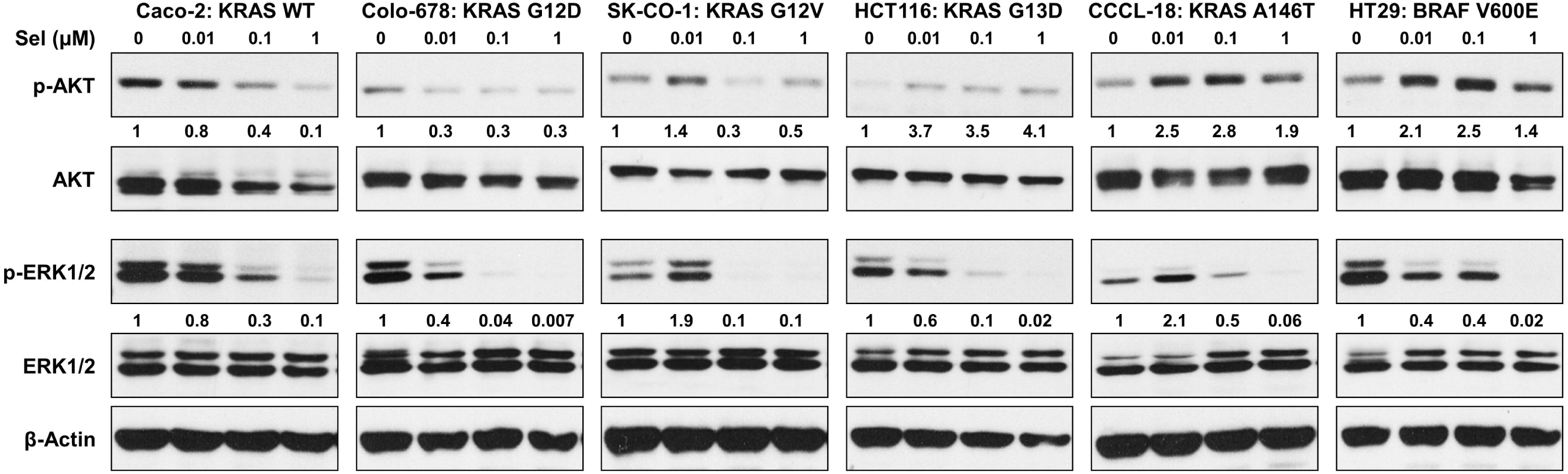
Inhibition of MEK leads to differential responses in RAS signaling in CRC cell lines. A panel of CRC cell lines including Caco-2 (KRAS WT), Colo-678 (KRAS G12D), SK-CO-1 (KRAS G12V), HCT116 (KRAS G13D), CCCL-18 (KRAS A146T), and HT29 (BRAF V600E) were treated with Sel and proteins were subject to Western blotting assay for assessment of phosphorylation of ERK and AKT.

**Figure 5.**
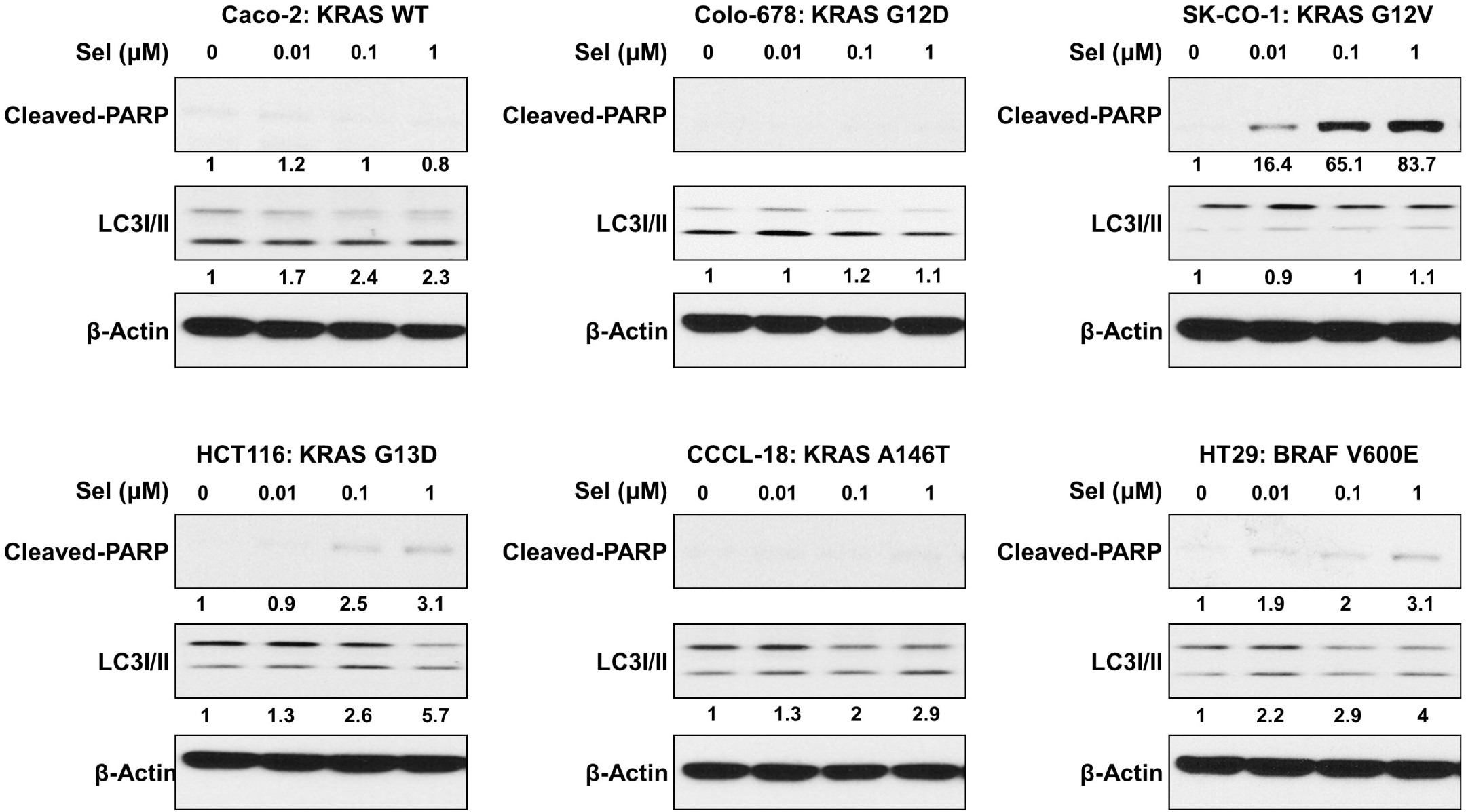
Inhibition of MEK modulates cell apoptosis and autophagy in a RAS allele specific manner in CRC cell lines. A panel of CRC cell lines including Caco-2 (KRAS WT), Colo-678 (KRAS G12D), SK-CO-1 (KRAS G12V), HCT116 (KRAS G13D), CCCL-18 (KRAS A146T), and HT29 (BRAF V600E) were treated with Sel and cell apoptosis and autophagy were examined as indicated by cleaved-PRAP and LC3 by Western blotting assay.

### Combinatorial approach of ALS and MEK inhibitor regulates cell apoptosis and autophagy in a RAS allele specific manner

At last, we examined the effects of combinatorial inhibition of AURAK and MEK via ALS and Sel on RAS signaling pathways and cell apoptosis and autophagy. We treated Caco-2, Colo-678, SK-CO-1, HCT116, CCCL-18, and HT29 cells with Sel (0.1 µM) for 24 hr in addition of ALS at 0.1, 1, and 5 µM for 24 hr. The RAS signaling output was evaluated via p-Akt and p-Erk and cell apoptosis and autophagy were examined via cleaved-PARP and LC3II/I. As shown in Figure 6, ALS counteracted the inhibitory effect of Sel on PI3K/Akt signaling pathway in Caco-2, HCT1116, CCCL-18, and HT29 cells, with increase in the level of p-Akt, whilst ALS enhanced the suppressive effect of Sel on PI3K/Akt signaling pathway in Colo-678 and SK-CO-1 cells, with further decline in the level of p-Akt. On the other hand, ALS strengthened the inhibitory effect of Sel on MAPK signaling pathway in Colo-678, SK-CO-1, and HT29 cells, with further suppression of Erk phosphorylation, whereas, ALS was counteractant to Sel in the rest of cell lines. As shown in Figure 7, ALS enhanced apoptotic effects in the presence of Sel in all tested cells except colo-678 cells. In particular, ALS enhanced the cleavage of PARP in wild-type KRAS expressing Caco-2 cells which did not occur in the presence of Sel alone. ALS enhanced autophagic effects in the presence of Sel in Caco-2, Colo-678, HCT116, and HT29 cells, but there was no alteration in SK-CO-1 and CCCL-18 cells. Taken together, combinatorial approach of ALS and Sel regulates cell apoptosis and autophagy in a RAS allele specific manner

**Figure 6.**
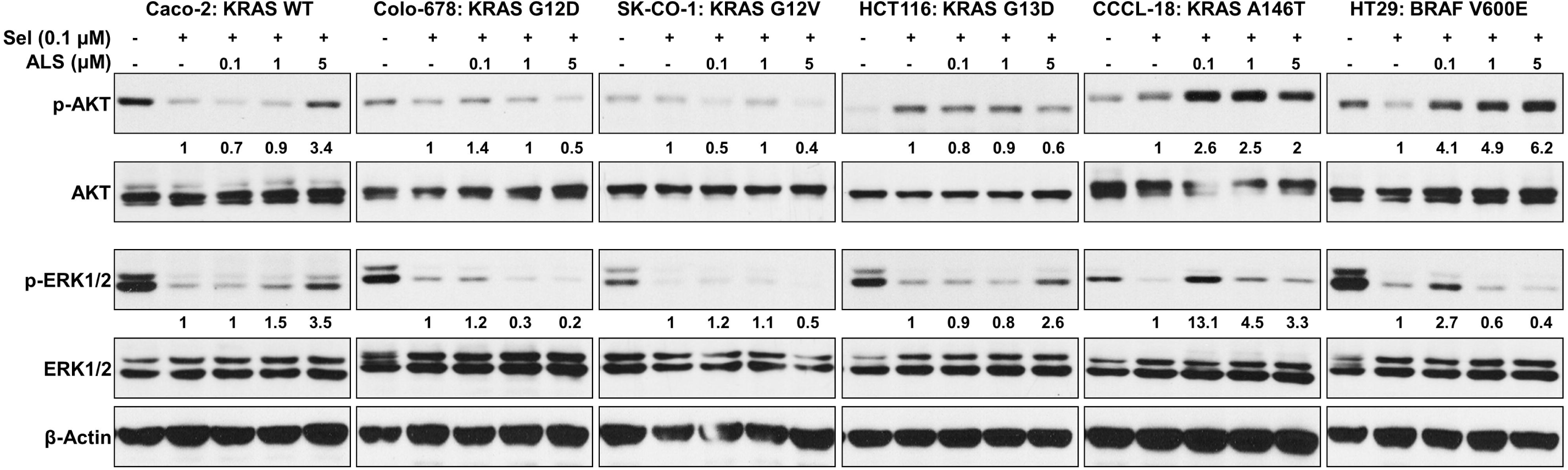
Combination of ALS and MEK inhibitor modulates PI3K/AKT and MAPK signaling in a RAS allele specific manner in CRC cell lines. A panel of CRC cell lines including Caco-2 (KRAS WT), Colo-678 (KRAS G12D), SK-CO-1 (KRAS G12V), HCT116 (KRAS G13D), CCCL-18 (KRAS A146T), and HT29 (BRAF V600E) were treated with ALS and Sel and proteins were subject to Western blotting assay for assessment of phosphorylation of ERK and AKT.

**Figure 7.**
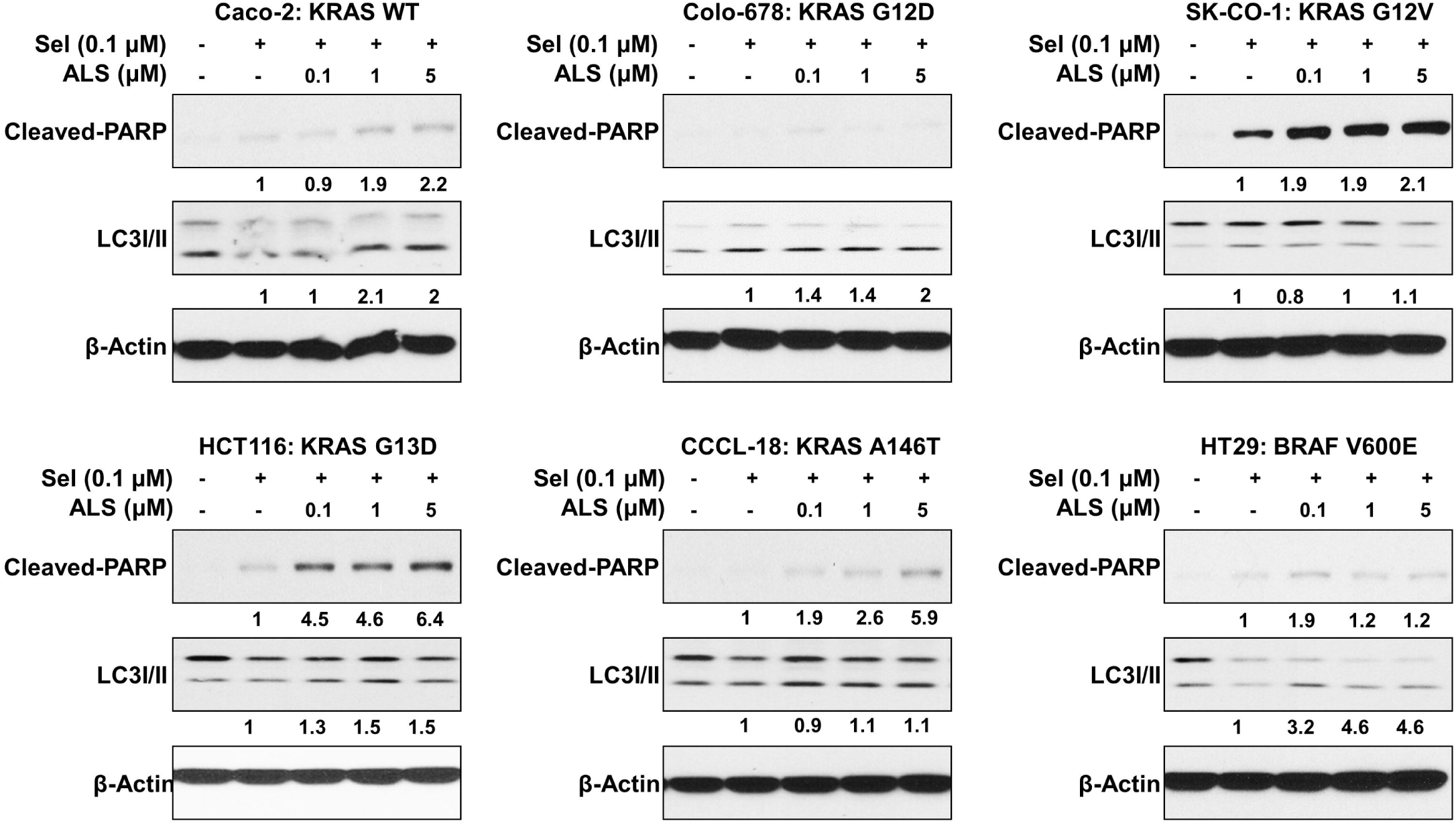
Combination of ALS and MEK inhibitor regulates cell apoptosis and autophagy in a RAS allele specific manner. A panel of CRC cell lines including Caco-2 (KRAS WT), Colo-678 (KRAS G12D), SK-CO-1 (KRAS G12V), HCT116 (KRAS G13D), CCCL-18 (KRAS A146T), and HT29 (BRAF V600E) were treated with ALS and Sel and cell apoptosis and autophagy were examined as indicated by cleaved-PRAP and LC3 by Western blotting assay.

### Combination of ALS and MEK inhibitor displays an enhanced cell growth inhibitory effect

At last, given the commonly mutated KRAS in CRC, we engineered Flp-In T-REx cells to express KRAS WT and G12D along with mCherry to assess the effects of ALS and Sel on cell growth alone or in combination. In a dose escalation assay, both monotherapies with ALS and Sel exerted stronger cell growth inhibitory effect in the context of KRAS G12D, relative to KRAS WT (Figure 8). Of note, the combination treatment with ALS and Sel displayed an enhanced cell growth inhibitory effect in KRAS G12D expressing cells, which is in consistence with the regulatory effect on signal output in KRAS G12D expressing CRC cell line. Collectively, it shows dual inhibition of AURKA and MEK would generate enhanced effect.

**Figure 8.**
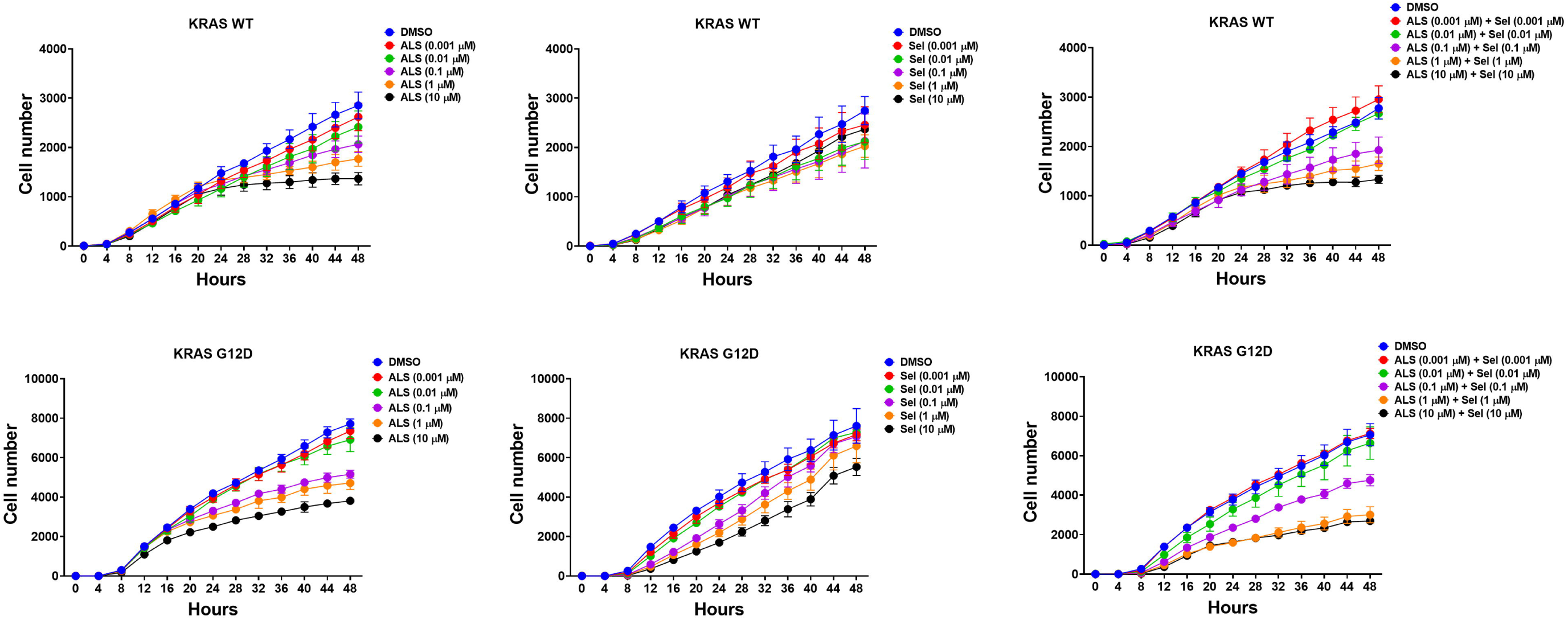
Dual inhibition of AURKA and MEK exerts improved cell growth inhibitor effect. Cell growth was examined in engineered Flp-In T-REx cells expressing KRAS WT and G12D along with mCherry in a dose escalation study. Cells were treated with ALS or Sel alone or in combination and the cell growth was monitored in real-time using IncuCyte.

## Discussion

Recently, accumulating evidence shows that RAS allele specific approach for cancer monotherapy or combination treatment resulting in encouraging outcome in preclinical and clinical settings. RAS driven CRC causes high mortality and morbidity [24], which needs novel therapeutic strategy associated with increased beneficial effects but decreased side effects. Given a large body of evidence showing that Aurora kinase is promising cancer therapeutic target [25-32], selective inhibition of this target in a RAS allele specific manner would produce profound therapeutic effect for cancer treatment. In the present study, a RAS allele specific regulatory effects on RAS signaling pathway of ALS was observed in a panel of CRC cell lines expressing different RAS mutants. We have found that ALS alone or in combination of MEK inhibitor exerts differential effects on PI3K/Akt and MAPK signaling pathway, and differentially induced cell apoptosis, autophagy, and cell growth.

The RAS subfamily consists of KRAS, NRAS, and HRAS, all of which are mutated in human cancers [33]. These isoforms demonstrate a high degree of sequence identity/similarity, except at the C-terminus. Nevertheless, they are mutated in cancer in a non-random distribution, suggesting context-dependent differences in biological function. NRAS mutations are most common in malignant melanoma, hematopoietic malignancies, and thyroid cancer while HRAS mutations are most common in head and neck and bladder cancers [33]. KRAS mutations are most common overall, occurring in up to 22% of all human cancers and predominate in lung cancer, CRC, and pancreatic cancers [33].

KRAS mutations typically occur in exons 2 and 3, at codons 12, 13, and 61 [33, 34]. Epidemiological and prospective clinical studies show that cancers expressing different mutant forms of KRAS exhibit distinct clinical behaviors [34, 35]. These differences likely arise from rewiring of signal transduction networks in a RAS mutation-dependent manner, which is inspiring studies on the RAS context dependency in signaling output. For example, a comparison of KRAS G12C, G12V, and G12D mutations in patient-derived NSCLC cell lines, demonstrate activation of MAPK and PI3K/AKT in G12D lines, whereas G12C and G12V show little PI3K/AKT signaling and prominent RAL activation [36]. Other studies have shown similar RAS allele-specific rewiring [37]. While these studies are critical for establishing that isoform or allele-specific effects occur, the mechanisms and translational implications have not been explored. Given this lack of understanding, RAS allele-specific biology is currently a major focus and RAS context dependency in monotherapy or combination therapy of other key nodes inhibitors with RAS signaling pathway inhibitors was a dominant topic in the RAS research community.

In conclusion, our findings unveil a therapeutically exploitable role for AURKA inhibition and RAS signaling modulation in the effort of CRC treatment in a RAS allele specific manner. Given that *KRAS* is the most common oncogene in human cancers, including CRC, novel therapeutic strategies are urgently needed. Not only may target a single kinase, but may also be therapeutically effective by harnessing oncogenic KRAS with an allele specific approach.

## Conflict of interest

The authors declare that there is no conflict of interest in this work.

## Acknowledgments

This work was supported by the Medical Science and Technology Research Project of Foshan City (2017AB003533), Medical Science and Technology Research Foundation of Guangdong Province (A2018150), Scientific Research Start Plan of Shunde Hospital, Southern Medical University (SRSP2018002).

## Notes

### Competing Interest Statement

The authors have declared no competing interest.

